# Ancient IL-6-STAT5 signaling orchestrates inflammation in jawless vertebrates

**DOI:** 10.64898/2026.04.02.716188

**Authors:** Francisco Fontenla-Iglesias, Jean-Louis Boulay, Jonathan P. Rast, Masayuki Hirano, Byron B. Au-Yeung, Jesús Lamas, Weiming Li, Thomas Boehm, Max D. Cooper, Sabyasachi Das

## Abstract

Interleukin-6 (IL-6) is a central regulator of vertebrate immunity, yet its existence in jawless vertebrates has remained obscure because of extreme sequence divergence. The extant jawless vertebrates (lampreys and hagfish), which deploy variable lymphocyte receptors (VLRs) instead of immunoglobulins and T cell receptors, provide a unique window into the earliest interface between cytokines and adaptive immunity. Here, we identify *IL-6-like* genes in jawless vertebrates by combining computational structural comparisons, phylogenetic reconstruction, and conserved local and long-range synteny analyses. The encoded proteins adopt the canonical four-helix bundle characteristic of immune-related IL-6 family cytokines and align topologically with mammalian IL-6. In sea lamprey, three *IL-6* paralogs show distinct leukocyte and tissue expression patterns and are differentially induced by pathogen-associated molecular patterns and skin injury, indicating early functional diversification of IL-6-mediated inflammatory responses. Stimulation of myeloid peritoneal leukocytes with recombinant lamprey IL-6 induces STAT5 phosphorylation and rapid upregulation of *SOCS1/3* genes, consistent with an IL-6/STAT5/SOCS regulatory axis. These findings extend the repertoire of immune-related four-helix bundle cytokines to jawless vertebrates and indicate that IL-6-dependent inflammatory programs were already in place before the divergence of VLR-based and Ig/TCR-based adaptive immune systems in vertebrates.

## Introduction

Approximately 500 million years ago, two fundamentally distinct forms of recombinatorial adaptive immunity (AIS) emerged in the earliest vertebrates, representing a major evolutionary innovation in host defense (Boehm 2025; Boehm et al. 2018; Cooper and Alder 2006; Flajnik and Kasahara 2010; Pancer et al. 2004). In jawed vertebrates, AIS is mediated by T and B lymphocytes that recognize antigens through immunoglobulin-domain receptors generated by V(D)J recombination and diversified via somatic hypermutation. By contrast, extant jawless vertebrates, the lampreys and hagfishes, lack these canonical jawed vertebrate AIS components and instead deploy variable lymphocyte receptors (VLRs) expressed by B-like and T-like lymphocytes. VLRs are assembled through the stepwise incorporation of diverse leucine-rich-repeat (LRR) genomic donor cassettes, providing an alternative yet equally complex dual-arm immune architecture. This parallel T- and B-cell framework likely arose in the last common ancestor of vertebrates and has been conserved in jawed and jawless vertebrates.

Among vertebrate cytokines, interleukin-6 (IL-6) is a paradigmatic pleiotropic factor that orchestrates immune regulation, inflammation, hematopoiesis, and tissue repair (Kishimoto and Ishizaka 1975) (Hirano 2020; Kang et al. 2020; Tanaka et al. 2014). Originally identified as a T-cell-derived B-cell stimulatory factor (Kishimoto and Ishizaka 1975), IL-6 is a four-helix-bundle cytokine that belongs to the wider IL-6 family, which also includes IL-11, ciliary neurotrophic factor (CNTF), leukemia inhibitory factor (LIF), oncostatin M (OSM), cardiotrophin (CT), cardiotrophin-like cytokine (CLC), and IL-27, among others (Rose-John 2018). Members of this family signal through receptor complexes that share the ubiquitously expressed transducer gp130 and activate canonical intracellular pathways such as JAK/STAT or MAPK (Murakami et al. 2019). In mammals, IL-6 is produced by a broad range of cell types, including macrophages, T and B lymphocytes, fibroblasts, and endothelial cells, among others (Kishimoto 2005) and its expression is induced by diverse pro-inflammatory cues such as IL-1β, TNF-α, microbial pathogen-associated molecular patterns (PAMPs), and tissue injury (Hirano 2020; Kang et al. 2019; Tanaka et al. 2014). Engagement of its heteromeric receptor, composed of the non-signaling IL-6Rα chain and the signal-transducing subunit gp130, triggers JAK1/2-mediated phosphorylation of gp130, followed by recruitment, phosphorylation, and dimerization of the signal transducer and activator of transcription 3 (STAT3), and its subsequent nuclear translocation to activate transcriptional programs that regulate cell proliferation, differentiation, and inflammation (Heinrich et al. 1998). To prevent chronic inflammation and autoimmunity, IL-6 signaling is tightly controlled by negative feedback loops mediated by suppressor of cytokine signaling (SOCS) proteins, particularly SOCS1 and SOCS3 (Tanaka et al. 2014).

Despite their central roles in immune regulation, the evolutionary histories of interleukins remain difficult to reconstruct because of extensive sequence divergence across vertebrate lineages, shaped in part by ongoing host-pathogen coevolution (Brocker et al. 2010). This divergence undermines the performance of classical homology-based approaches such as BLAST (Altschul et al., 1990), in which pairwise nucleotide or amino acid sequence identity is often too low to confidently infer orthology. IL-6 orthologs frequently exhibit very limited primary sequence similarity, particularly between bony and cartilaginous fishes, complicating efforts to establish evolutionary continuity. This challenge is even more pronounced in jawless vertebrates, where the absence of recognizable IL-6 orthologs has long impeded both functional and evolutionary analyses.

In this study, we identify and functionally characterize *IL-6* candidate genes in lamprey through an integrated framework combining large language model (LLM)-based structural modeling, phylogenetic inference, and conserved synteny analysis. We show that *IL-6* expression is induced by canonical PAMP and skin injury, recapitulating the transcriptional responsiveness described in mammals. We further show that IL-6 signaling is transmitted through STAT5 phosphorylation, resulting in SOCS induction and pointing to an evolutionarily conserved signaling mechanism. These findings support the existence of an IL-6-like cytokine axis in the last common ancestor of jawed and jawless vertebrates and indicate that key features of IL-6-mediated inflammatory signaling were already in place before the split of these two major vertebrate lineages.

## Results

### Identification of IL-6 orthologs in lampreys and hagfish

To identify IL-6 homologs in jawless vertebrates, we first conducted sequence similarity searches using BLAST against available nucleotide and protein datasets from the sea lamprey (*Petromyzon marinus*) and Atlantic hagfish (*Myxine glutinosa*), using IL-6 sequences from representative jawed vertebrates as queries. No significant hits were recovered, consistent with the extensive primary-sequence divergence characteristic of IL-6 cytokines. We therefore implemented a structure-guided discovery pipeline grounded on the defining features of IL-6-type cytokines: (i) a predicted protein length of 150-280 amino acids; (ii) an N-terminal signal peptide; (iii) a canonical four-helix bundle topology predicted by AlphaFold (Jumper et al. 2021) or ESMFold (Lin et al. 2023); and (iv) high structural similarity to known IL-6 protein in jawed vertebrates as assessed by FoldSeek (van Kempen et al. 2024). This integrative approach identified a single candidate in hagfish and four IL-6 family-like candidates in sea lamprey (Figs. 1, S1). One of the four sea lamprey candidates, located on chromosome 4, was identified as a cardiotrophin-2 (CT-2) homolog on the basis of reciprocal BLAST, phylogenetic analysis, and the absence of the canonical cysteine configuration that typifies IL-6 proteins (Figs. 1, S1). Notably, *CT-2* is present as a pseudogene in humans (Derouet et al. 2004). In mammals, IL-6 proteins typically contain four conserved cysteine residues, which form two intramolecular disulfide bonds (Northemann et al. 1989; Snouwaert et al. 1991). This canonical configuration is retained in the lamprey IL-6 candidate sequences. In contrast, the hagfish IL-6 candidate conserves only two cysteine residues (Fig. S1), a feature also observed in some bony fish IL-6 orthologs (Bird et al. 2005; Eggestol et al. 2020).

**Figure 1.**
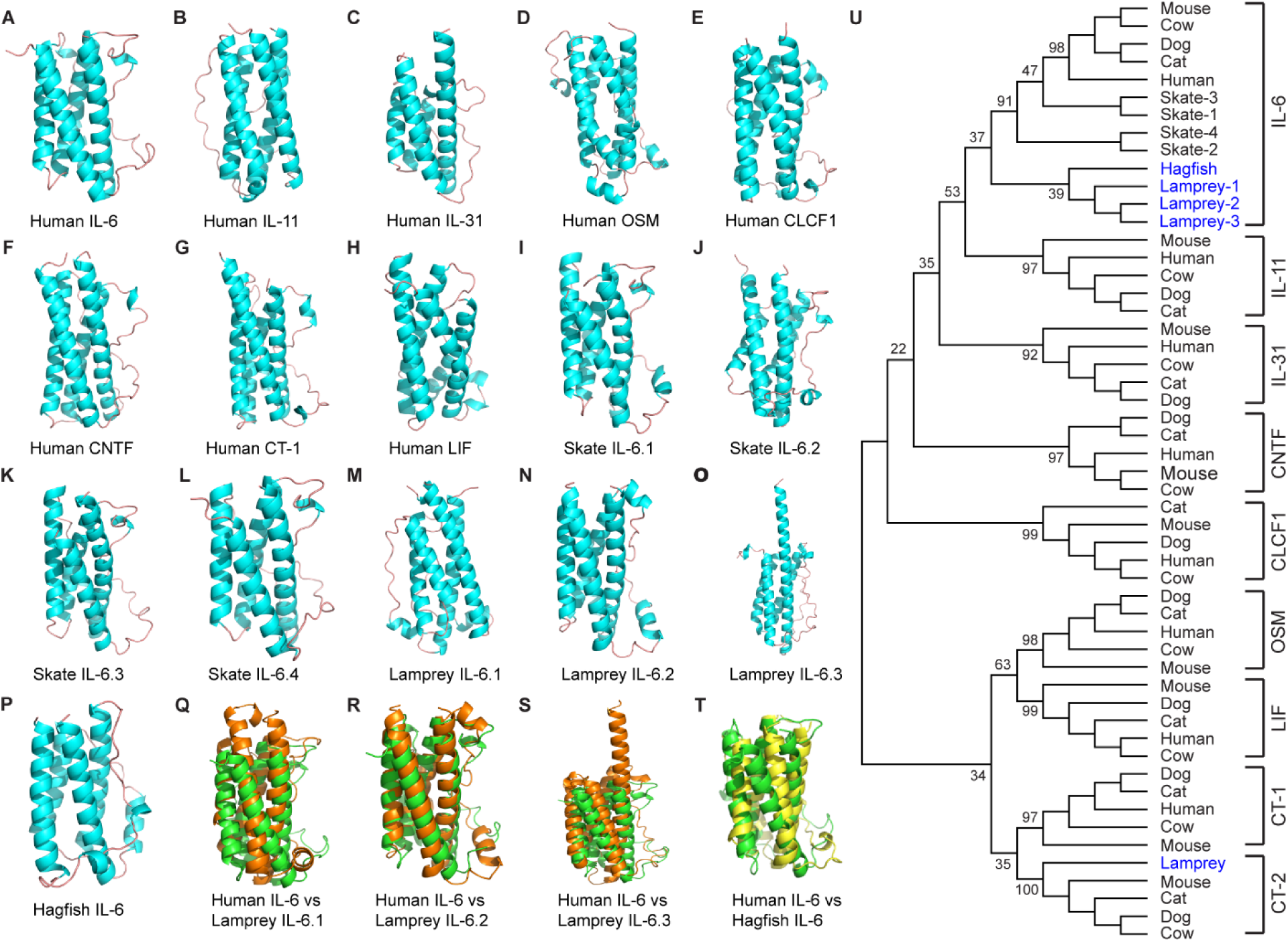
Conserved structural features of IL-6 homologs in jawless vertebrates. (A-H) Predicted three-dimensional structures of representative human IL-6 family cytokines: IL-6, IL-11, IL-31, oncostatin M (OSM), cardiotrophin-like cytokine factor 1 (CLCF1), ciliary neurotrophic factor (CNTF), cardiotrophin-1 (CT-1), and leukemia inhibitory factor (LIF). Despite differences amino acid composition, all adopt the canonical four-helix bundle. (I-P) Predicted tertiary structures of IL-6 sequences from thorny skate (*Amblyraja radiata*) and putative IL-6 homologs from sea lamprey (*Petromyzon marinus*; three sequences designated IL-6.1-IL-6.3) and Atlantic hagfish (*Myxine glutinosa*), each exhibiting a conserved four-helix topology with an IL-6-like domain. (Q-T) Structural alignments of human IL-6 (green color) with the lamprey and hagfish orthologs show strong topological conservation despite substantial sequence divergence. (u) Phylogenetic analysis places the lamprey and hagfish putative IL-6 homologs closer to mammalian IL-6 than to other IL-6 family cytokines; bootstrap support values are indicated at internal nodes.

Like human IL-6 family members (Fig. 1A–H) and the four IL-6 sequences identified in thorny skate (Fig. 1I–L), all lamprey and hagfish candidates displayed the characteristic four-helix bundle architecture (Fig. 1M–P) and resembled human IL-6 structurally (Fig. 1Q–T). The three sea lamprey IL-6 candidates were also structurally alignable with the Atlantic hagfish IL-6 candidate (Fig. S2). However, the third sea lamprey sequence, which contains an extended α-helical region, showed weaker structural alignment with both the hagfish IL-6 candidate (Fig. S2) and human IL-6 (Fig. 1S). In phylogenetic analyses including representative mammalian IL-6 family members (Fig. 1U), the jawless vertebrate sequences clustered within the canonical IL-6 clade, although this placement was weakly supported (bootstrap support = 37%). Thus, the phylogenetic signal alone is insufficient to confidently establish orthology. Nevertheless, the concordance between structural similarity and phylogenetic placement provides preliminary support for the interpretation that these sequences represent IL-6 orthologs rather than other members of the broader IL-6 cytokine family.

Genomic context analyses further support the assignment of these genes as *IL-6* orthologs (Fig. 2). We first inspected the immediate genomic neighborhood of *IL-6* using the NCBI Genome Data Viewer and found that *IL-6* lies directly adjacent to *TOMM7* in human, chicken, spotted gar, and thorny skate, with the two genes arranged in opposite transcriptional orientations (Fig. S3). The hagfish *IL-6* gene shows the same *TOMM7*-*IL-6* head-to-head configuration (Fig. S3), indicating retention of this ancestral microsyntenic relationship. In sea lamprey, by contrast, *TOMM7* is located on a short scaffold that is physically separated from all three *IL-6* loci, suggesting disruption of local gene order (Fig. 2A).

**Figure 2.**
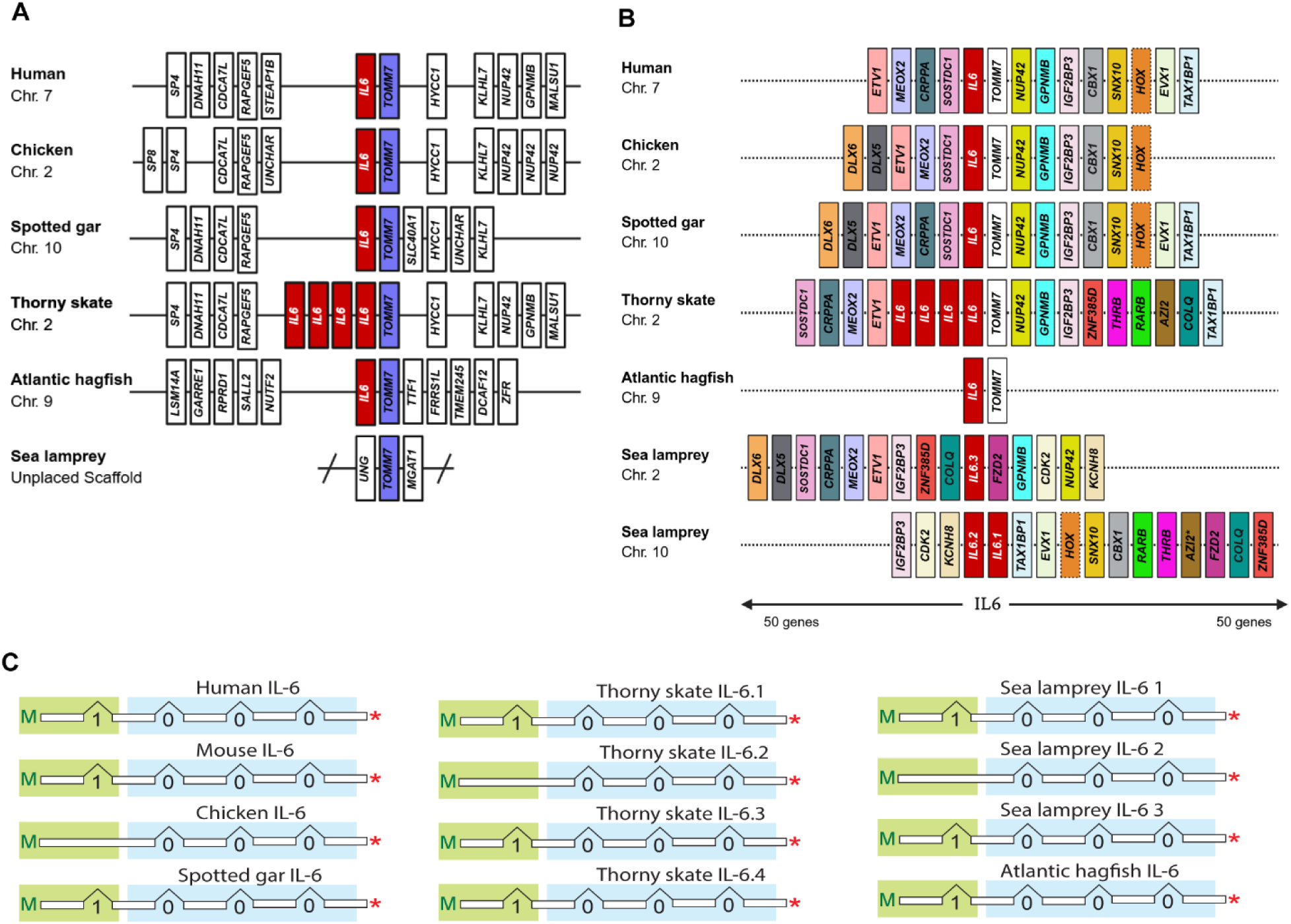
Genomic organization and synteny conservation of *IL-6* loci across cyclostomes and gnathostomes. (A) Analysis of the *TOMM7*-*IL-6* microsyntenic relationships across representative vertebrates, including *Homo sapiens* (Human), *Gallus gallus* (Chicken), *Lepisosteus oculatus* (Spotted Gar), *Amblyraja radiata* (Thorny skate), *Myxine glutinosa* (Atlantic hagfish), and *Petromyzon marinus* (Sea lamprey). (B) Comparative analysis of the genomic neighborhoods flanking IL6 across vertebrates. Fifty genes upstream and downstream of *IL-6* in *H. sapiens* (Chr. 7) were used as a reference window, and orthologs were mapped in *G. gallus* (Chr. 2), *L. oculatus* (Chr. 10), *A. radiata* (Chr. 2), *M. glutinosa* (Chr. 9), and *P. marinus* (Chr. 2 and Chr. 10). Colored blocks denote conserved orthologs. In jawed vertebrates, *IL-6* resides within a deeply conserved block containing *ETV1*, *MEOX2*, *CRPPA*, *SOSTDC1*, *TWIST2*, *TOMM7*, *NUP42*, *GPNMB*, *IGF2BP3*, *CBX1*, *SNX10*, *EVX1*, and *TAX1BP1*. In hagfish, *IL-6* retains synteny only with *TOMM7*, while in sea lamprey three *IL-6* paralogs are distributed across two chromosomes, each retaining subsets of the conserved neighborhood. The orange dashed box highlights a variable cluster of Hox-related genes. (C) Conservation of exon-intron architecture of *IL-6* genes. Schematic representation of exon-intron organization between the start codon (M) and the stop codon (*) in representative *IL-6* genes, illustrating conserved intron phasing within the signal peptide (green box) and *IL-6* domains (blue box). Accession numbers are provided in the Supplementary Table S1.

To extend this analysis, we expanded the analysis to the 50 protein-coding genes upstream and downstream of each *IL-6* locus in all species examined (Fig. 2B). This broader comparison revealed that many genes that flank *IL-6* in gnathostomes, including *ETV1*, *MEOX2*, *CRPPA*, *SOSTDC1*, *TWIST2*, *GPNMB*, *IGF2BP3*, *CBX1*, *SNX10*, *EVX1*, and *TAX1BP1*, are retained in the vicinity of the lamprey *IL-6* loci but partitioned between sea lamprey chromosomes 10 and 2. This partitioning of conserved neighboring genes indicates that the gnathostome *IL-6* locus likely corresponds to a combination of these two lamprey genomic regions. Such an arrangement is consistent with a segmental duplication of an ancestral chromosomal block containing *IL-6*, rather than duplication of the *IL-6* gene alone. *IL-6.1* and *IL-6.2* occur as tandem paralogs on chromosome 10, whereas *IL-6.3* is located on chromosome 2. Several flanking genes, such as *ZNF385D*, *KCNH8*, and *COLQ*, are themselves duplicated and differentially associated with the *IL-6.1*/*IL-6.2* and *IL-6.3* regions, further supporting the paralogous relationship between these loci (Fig. 2B). We also examined the *IL-6* loci in a distantly related lamprey species, the pouched lamprey (*Geotria australis*), which diverged from the sea lamprey lineage approximately 250 million years ago (Kuraku and Kuratani 2006). As in sea lamprey, *G*. *australis* possesses *IL-6.1* and *IL-6.2* in tandem on the same chromosome or scaffold, whereas *IL-6.3* resides at a distinct genomic locus (Table S1). These findings indicate that the segmental duplication giving rise to the tandem *IL-6.1*/*IL-6.2* arrangement, followed by the chromosomal rearrangement that separated *IL-6.3*, occurred early in lamprey evolution. This split conservation pattern is consistent with the extensive segmental duplication and chromosomal reshuffling reported for agnathan genomes (Smith et al., 2018; Marlétaz et al., 2024; Yu et al., 2024), which has eroded local gene order while preserving long-range syntenic relationships to the gnathostome *IL-6* region.

Consistently, the macrosynteny chord diagram of human and sea lamprey genes shows that both sea lamprey chromosomes 10 and 2 share broad syntenic correspondence with human chromosome 7, which harbors *IL-6* (Fig. S4). These relationships indicate that the *IL-6*-containing genomic segments on the two lamprey chromosomes are paralogous to one another and co-orthologous to the corresponding segment of human chromosome 7, reflecting duplication and/or rearrangement of an ancestral *IL-6*-bearing chromosome early in the lamprey lineage.

In addition to chromosomal localization, gene structure provides an independent line of evidence for evolutionary conservation. We therefore compared the exon-intron organization and intron phasing of IL-6 coding-region introns across representative jawed vertebrates, lamprey, and hagfish (Fig. 2C). In most species examined, *IL-6* genes are composed of five exons and four introns with conserved intron phases, including one intron in the signal peptide-coding region and three in the IL6 domain. A notable exception was observed in *IL-6.2* of sea lamprey and in thorny skate, in which the intron within the signal peptide region has been lost. Overall, this conserved exon-intron architecture supports a shared evolutionary origin of *IL-6* genes.

### *IL-6* genes display discrete leukocyte and tissue expression in sea lamprey

To define the cellular expression landscape of the three *IL-6* genes, we sorted ammocoete-stage white blood cells using monoclonal antibodies against VLRA, VLRB, VLRC, monocytes, and granulocytes (Guo et al. 2009; Han et al. 2015; Hirano et al. 2013). Quantitative RT-PCR revealed a sharply partitioned expression pattern for the three *IL-6* paralogs in sea lamprey (Fig. 3). *IL-6.1* was enriched in monocytes, B-like (VLRB⁺) lymphocytes, and triple-negative (VLRA⁻/VLRB⁻/VLRC⁻) lymphocytes. *IL-6.2* transcripts predominated in granulocytes and triple-negative lymphocytes, whereas *IL-6.3* expression was confined to granulocytes and monocytes. All three paralogs were barely detectable in VLRA⁺ or VLRC⁺ T-like cell populations (Fig. 3A); owing to the lack of isotype-specific antoiboides, the expression pattern of the paralogs could be assessed in VLRD-, VLRE-, and VLRF-expressing cells.

**Figure 3.**
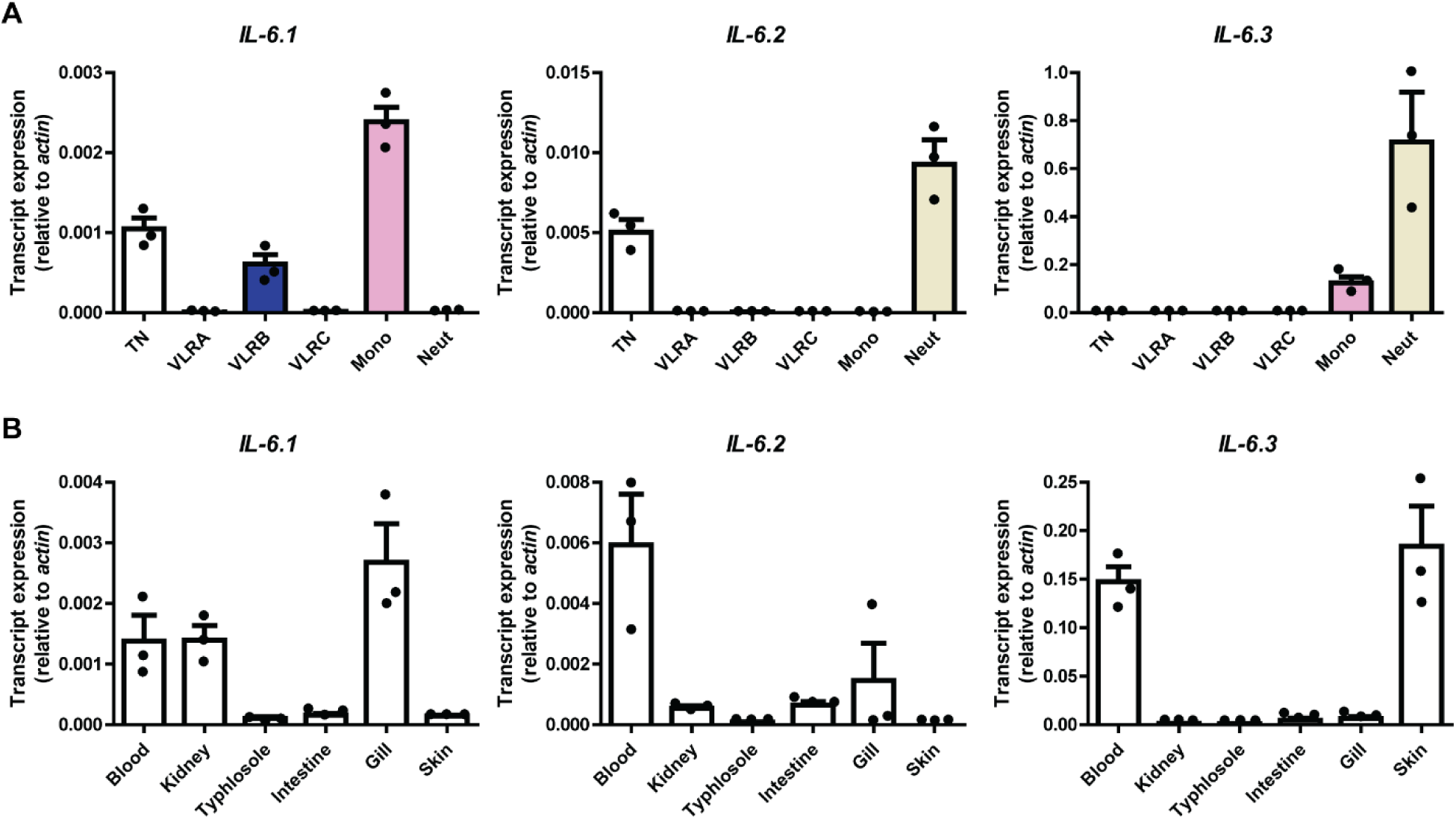
Divergent cellular and tissue expression landscapes of lamprey *IL-6* paralogs. (A) Expression of the *IL-6* gene in different lymphocyte populations in lamprey larvae. TN represents the triple-negative (VLRA^−^/VLRB^−^/VLRC^−^) lymphocyte population. Mono: Monocyte, Neut: Neutrophil. Bars indicate the standard error of the mean in each experiment (n=3). (B) Tissue expression profiles for *IL-6* in lamprey larvae. Transcripts are analyzed using real-time RT-PCR, with β-actin serving as a normalization control for both cellular and tissue distribution analyses. Bars indicate the standard error of the mean for three animals in each experiment. The corresponding data points were shown as dot plots. Bars represent mean ± SD.

Basal tissue profiling of larval immune-associated organs further underscored these distinctions (Fig. 3B). *IL-6.1* transcripts were highest in gill, moderate in blood and kidney, and minimal in typhlosole, intestine, and skin. *IL-6.2* was expressed primarily in blood, with lower abundance in kidney, typhlosole, intestine, and gill, and was undetectable in skin. In contrast, *IL-6.3* was strongly expressed in blood and skin and at lower levels in other tissues. Within the context of these measurements, these data suggest that the individual lamprey *IL-6* genes have evolved distinct leukocyte and tissue-specific expression patterns, consistent with functional specialization in lamprey immunity.

### Lamprey *IL-6* genes are differentially induced during immune stimulation and tissue injury

Ammocoete-stage sea lampreys were injected with flagellin or phosphate-buffered saline (PBS), and transcript levels of the three *IL-6* genes were quantified by RT-qPCR in the typhlosole/intestine at 2, 8, and 24 hours post-injection. All three *IL-6* paralogs were induced by flagellin, with *IL-6.1* showing a strong transient peak at 2 hours and declining thereafter, *IL-6.2* displaying a broader induction, and *IL-6.3* reaching its maximal expression at 8 hours (Fig. 4A-C). Expression of all three genes returned to near-basal levels by 24 hours. The inflammatory marker *IL-8* exhibited a similar early induction pattern (Fig. 4D). In a separate experiment, mechanical wounding of the skin for 2 hours resulted in upregulation of *IL-6.2* and *IL-6.3*, whereas *IL-6.1* expression remained unchanged (Fig. 4E-G). *IL-8* was also strongly induced, consistent with a local inflammatory response and likely leukocyte recruitment to the wounded area (Fig. 4H). Together, these findings suggest that *IL-6.1* responds more selectively to microbial PAMP, whereas *IL-6.2* and *IL-6.3* are activated more broadly under both microbial challenge and wounding-associated inflammatory conditions. Because microbial products from the skin surface or aquarium environment cannot be excluded in this setting, however, the wounding response should not be interpreted as reflecting strictly sterile inflammation. Rather, these data support differential responsiveness of lamprey *IL-6* paralogs to distinct inflammatory contexts, consistent with early functional diversification within this cytokine family.

**Figure 4.**
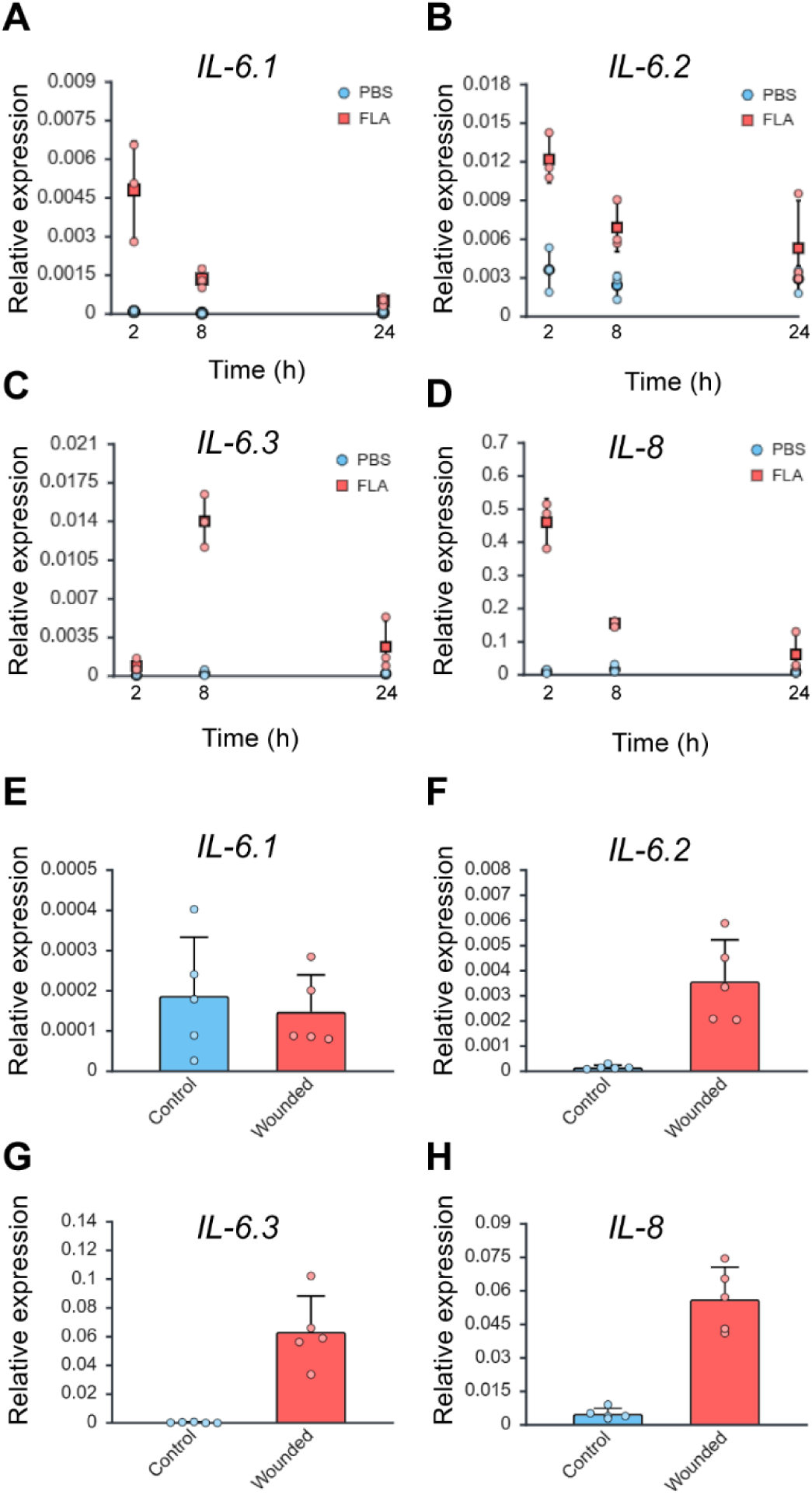
*IL-6* genes show distinct transcriptional responses to PAMP stimulation and tissue injury. (A-D) RT-qPCR analysis of *IL-6.1*, *IL-6.2*, *IL-6.3*, and *IL-8* expression in the typhlosole/intestine of sea lamprey larvae 2, 8, and 24 h after intraperitoneal stimulation with *Salmonella typhimurium*-derived flagellin. Control animals received PBS injections; n = 3 per condition. (E-H) RT-qPCR analysis of the same genes in the skin 2 h after injury induced by syringe puncture; uninjured animals served as controls; n = 6. Expression levels were normalized to *β-actin* and expressed as relative fold change compared with the corresponding control group. Bars represent mean ± SD.

### STAT5 phosphorylation is selectively induced by PmIL-6.1 and PmIL-6.2 in lamprey

To assess the biological activity of the three *IL-6* paralogs, we performed cytokine-stimulation assays using recombinant lamprey proteins produced in Chinese hamster ovary (CHO) cells. Because sufficient cell numbers could not be obtained from ammocoetes for in vitro assays, adult non-feeding sea lampreys were used as a source of inflammatory cells. To generate a reproducible population of responsive myeloid peritoneal exudate cells, we adapted a thioglycolate-induced peritonitis protocol originally developed in mice (Stewart et al. 1975) (Fig. 5A). Bulk RNA sequencing of unstimulated lamprey peritoneal cells indicated high levels for monocyte/macrophage-associated transcripts, including *GRN*, *CTSL*, *EFHD2*, *MMP2*, *PLA2G15* and *MARCO* (Fig. 5B), previously identified in monocytes and macrophages by single-cell RNA sequencing (Das et al. 2025; Huang et al. 2024).*VLR* transcripts were detected at lower levels overall in this cell population (Fig. 5C), although *VLRB* was more abundant than the other *VLR* genes examined. Together, these data suggest that the peritoneal exudate cell preparation contains a substantial fraction of myeloid-like cells.

**Figure 5.**
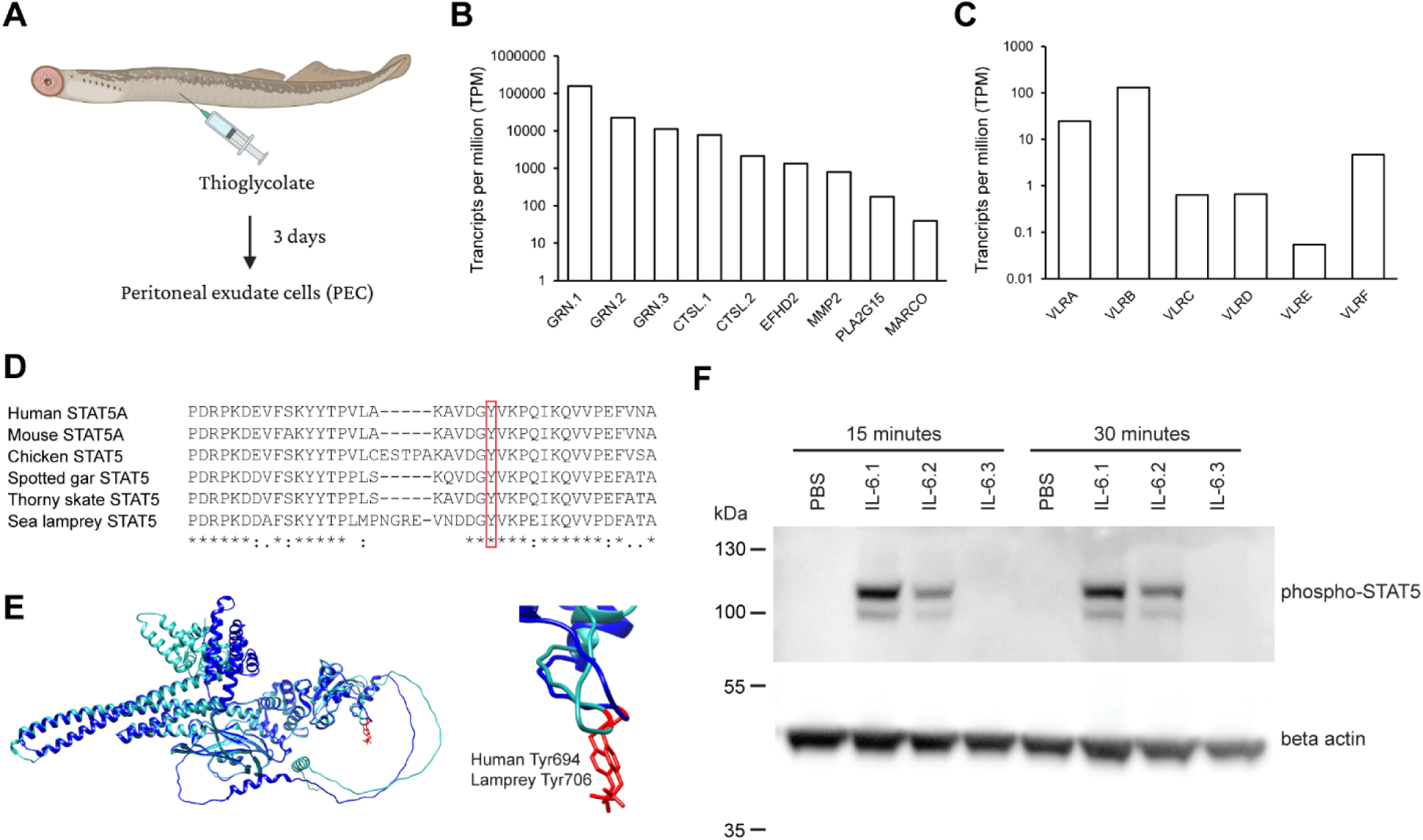
Lamprey IL-6 paralogs activate STAT5 signaling in peritoneal inflammatory cells. (A) Macrophage-like peritoneal exudate cells (PECs) were isolated from adult Petromyzon marinus three days after intraperitoneal thioglycolate injection. (B-C) Bulk RNA-seq profiling of PECs revealed robust expression of monocyte/macrophage-associated markers, including *GRN.1*, *GRN.2*, *GRN.3*, *CTSL.1*, *CTSL.2*, *FTH1*, *EFHD2*, *MMP2*, *PLA2G1*, and *MARCO* (B), together with low-to-moderate expression of *VLR* gene families (*VLRA*, *VLRB*, *VLRC*, *VLRD*, *VLRE*, *VLRF*). Transcript abundance is shown as TPM on a log scale. (D) Multiple sequence alignment of STAT5 orthologs from human, mouse, chicken, spotted gar, thorny skate, and sea lamprey identifies the conserved regulatory tyrosine phosphorylation site (red box). (E) Structural alignment of the AlphaFold3-predicted lamprey STAT5 model (light blue) with human STAT5B (deep blue), highlighting the O-phospho-L-tyrosine modification (red). Right, close-up of the conserved tyrosine residue positioned within a flexible loop region. (F) Whole-cell lysates from lamprey PECs stimulated *in vitro* with recombinant IL-6.1, IL-6.2, or IL-6.3 for 15 or 30 min were subjected to immunoblot analysis using antibodies to phosphorylated STAT5 and β-actin. Data shown are representative of three independent biological experiments. PBS served as the unstimulated control.

In jawed vertebrates, canonical IL-6 signaling proceeds via gp130-associated JAKs and activates STAT3 as the principal transcriptional effector, with additional activation of STAT1 (Heinrich et al. 2003) and, in specific cellular contexts, STAT5 (Piekorz et al. 1997; Tormo et al. 2012). In lampreys and hagfish, we identified two STAT1/3-like sequences, STAT1/3.1 and STAT1/3.2, that share substantial amino acid similarity, exhibit syntenic relationships with both STAT1 and STAT3, and cluster within the clade containing gnathostome STAT1 and STAT3 sequences in our phylogenetic analyses (Fig. S5). By contrast, we identified two STAT5 members (STAT5.1 and STAT5.2) in lampreys and hagfish that cluster with jawed vertebrate STAT5 orthologs with strong bootstrap support (Fig. S5). These findings are consistent with previous genomic analyses indicating that lampreys lack a clear STAT3 ortholog but retain a STAT5 ortholog (Boulay et al. 2022).

To probe downstream effectors of IL-6 signaling in lamprey, we leveraged the conservation of STAT activation motifs between mammals and agnathans (Fig. 5D). Reasoning that epitope conservation might permit antibody cross-reactivity, we tested commercial monoclonal antibodies against phosphorylated human STAT1 and STAT5 on lamprey leukocyte lysates. Upon treatment with pervanadate, a broad-spectrum tyrosine-phosphatase inhibitor that enforces global tyrosine phosphorylation, the anti-pSTAT5 antibody reproducibly detected two closely migrating bands at ∼100 kDa, consistent with the expected mass of STAT5 and with the presence two STAT5-like paralogs in lamprey (Fig. S6). In contrast, anti-pSTAT1 failed to yield a detectable signal, suggesting insufficient epitope conservation and precluding direct assessment of STAT1 phosphorylation. We therefore focused subsequent analyses on STAT5 (Fig. S6).

Peritoneal exudate cells were stimulated in vitro with recombinant PmIL-6.1, PmIL-6.2, or PmIL-6.3. Both PmIL-6.1 and PmIL-6.2 induced robust STAT5 phosphorylation detectable at 15 minutes, sustained at 30 minutes, whereas neither PmIL-6.3 nor unstimulated controls elicited STAT5 activation (Fig. 5F). This transient phosphorylation pattern is consistent with tightly regulated cytokine signaling and supports the interpretation that STAT5 participates in signaling downstream of PmIL-6.1 and PmIL-6.2 in lamprey. However, contributions from other STAT family members cannot be excluded, particularly given the antibody cross-reactivity limitations that prevented their evaluation in the present study.

### Recombinant IL6.1 and IL6.2 induces the upregulation of *SOCS* genes

SOCS proteins act as early-response negative regulators of JAK-STAT signaling, and in mammals, specific family members, particularly SOCS1 and SOCS3, are preferentially induced by IL-6(Sobah et al. 2021). Using sequence similarity, phylogenetic analysis, and conserved synteny with vertebrate SOCS family members, we identified three SOCS1/3-like genes (SOCS1/3.1, SOCS1/3.2, and SOCS1/3.3) and one SOCS2-like gene in lamprey and hagfish (Fig. 6A, Fig. S7). In phylogenetic trees, vertebrate SOCS2 grouped with CISH, whereas SOCS1 and SOCS3 formed a separate cluster, albeit with low support (Fig. 6A). Synteny analysis likewise did not resolve the relationships among the three *SOCS1/3*-like genes in jawless vertebrates, leaving their exact correspondence to vertebrate *SOCS1* and *SOCS3* uncertain (Fig. S7).

**Figure 6.**
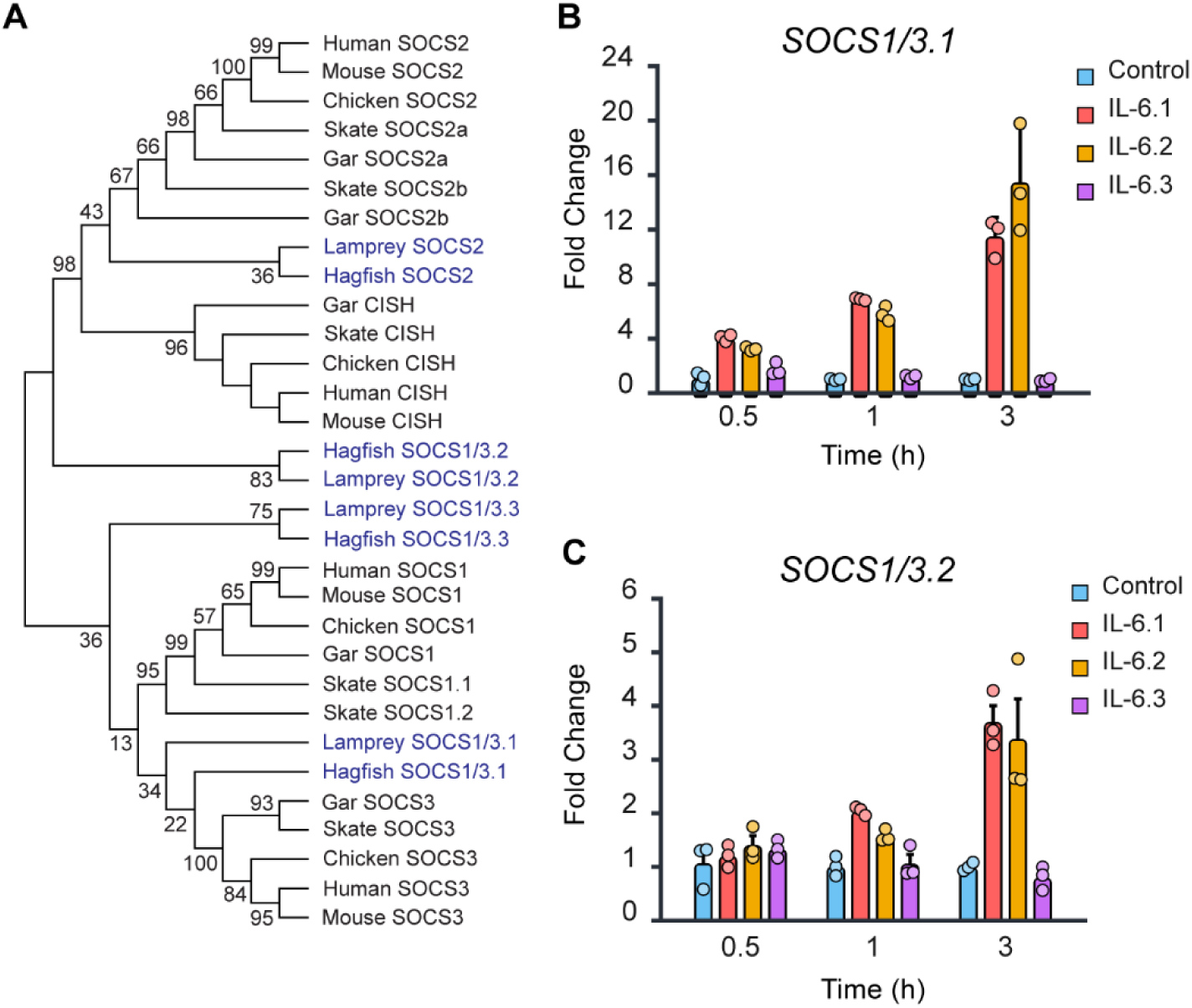
Lamprey IL-6 paralogs induce SOCS gene expression. (A) Phylogenetic analysis of SOCS1, SOCS2, SOCS3 and CISH family members of human, mouse, chicken, spotted gar, thorny skate, and sea lamprey. (B-C) RT-PCR analysis of *SOCS1/3.1* (B) and *SOCS1/3.2* (C) expression in lamprey PECs stimulated in *vitro* with IL-6.1, IL-6.2, or IL-6.3 recombinant proteins for 30 min, 1 h, and 3 h (n=3 wells per condition). Data are representative of two independent experiments. Expression values were normalized to β-actin and expressed as fold change relative to PBS controls. Bars represent mean ± SD.

We assessed the expression dynamics of the four *SOCS* genes in peritoneal exudate cells following *in vitro* stimulation with recombinant PmIL-6 paralogs. *SOCS1/3.1* was rapidly and strongly upregulated by PmIL-6.1 and PmIL-6.2, with transcript levels increasing at 30 minutes and reaching approximately 12-fold and 18-fold induction, respectively, at 3 hours post-stimulation. No changes were observed in cells stimulated with PmIL-6.3 or in unstimulated controls (Fig. 6B). In contrast, *SOCS1/3.2* exhibited a moderate response to PmIL-6.1 and PmIL-6.2, showing a gradual increase that peaked at ∼3.5-fold induction at 3 hours. Again, PmIL-6.3 and control conditions did not elicit detectable induction (Fig. 6C). Expression of the remaining *SOCS* genes (*SOCS1/3.3* and *SOCS2*) was too low under our experimental conditions to permit reliable quantification.

## Discussion

Although the conservation of interleukin receptor components across gnathostomes and cyclostomes suggests the presence of interleukin networks in jawless vertebrates (Boulay et al. 2022), interleukin orthologs have been difficult to identify outside jawed vertebrates. This difficulty reflects the fact that these signaling molecules are small proteins that exhibit extensive primary-sequence divergence even among jawed vertebrate classes, driven by strong host-pathogen co-evolution (Brocker et al. 2010), which often obscures detectable sequence similarity. Conservation of local synteny has been successfully used to pinpoint highly divergent interleukin genes (Dijkstra 2014; Fontenla-Iglesias et al. 2025; Holt et al. 2011). However, relying on synteny alone has important limitations in agnathans, because extensive segmental duplications and large-scale chromosomal rearrangements have reorganized local gene neighborhoods (Smith et al., 2018; Marlétaz et al., 2024; Yu et al., 2024). We recently overcame these limitations for the IL-1 family by using an integrated approach that combined computational three-dimensional structural conservation, conserved synteny, and functional assays to identify IL-1 orthologs across eumetazoans (Fontenla-Iglesias et al. 2025). Here, we apply a similar strategy to resolve the evolutionary history and function of IL-6 in jawless vertebrates.

IL-6 is widely recognized as an amplifier of acute and chronic inflammation, yet accumulating evidence shows it is a versatile regulator of physiology and disease (Tanaka et al. 2014). How and when this cytokine emerged during vertebrate evolution has remained unclear. By combining improved lamprey and hagfish genome assemblies with modern protein structure-prediction tools, we detected and modeled IL-6-like cytokines in jawless vertebrates, including three orthologs in lamprey and one in hagfish. Beyond establishing the presence of an IL-6/STAT5 module in jawless vertebrates, our data begin to outline how this axis is wired into the broader cytokine network and adaptive-like immunity. The biased expression of *IL-6.1* in monocytes, VLRB⁺ B-like lymphocytes, and triple-negative lymphocytes, together with its preferential induction by microbial PAMP, is consistent with a role in coupling myeloid sensing of infection to lymphocyte activation. By contrast, the broader responsiveness of *IL-6.2* and *IL-6.3* to both flagellin and tissue injury, and their enrichment in granulocytes and skin, point to a division of labor among distinct *IL-6* paralogs that preferentially support inflammatory and repair programs. This interpretation is further reinforced by recent findings showing that lamprey IL-6 receptors display distinct expression profiles across tissues and developmental stages, as well as dynamic regulation in response to environmental and metabolic changes (Gong et al. 2025).

At the level of downstream signaling, we found that STAT5 participates in the of IL-6-mediated signaling cascade in lampreys. Although STAT3 is the canonical mediator of IL-6 signaling in mammals, a clear STAT3 ortholog is absent in both lamprey and hagfish (Boulay et al. 2022). In jawed vertebrates, canonical IL-6 signaling predominantly engages STAT3 as the principal transcriptional effector, with additional activation of STAT1 (Heinrich et al. 2003) and, in specific cellular contexts, STAT5 (Piekorz et al. 1997; Tormo et al. 2012). Our findings nevertheless raise the possibility that STAT5 served as an ancestral downstream effector of IL-6 signaling in jawless vertebrates, preceding the clearer diversification of STAT1 and STAT3 and the signaling specialization that later emerged in jawed vertebrates. In this context, the development of reagents capable of detecting phosphorylation of lamprey STAT1/3-like proteins will be important for determining whether IL-6 can also engage these pathways. Accordingly, the ability of lamprey IL-6.1 and IL-6.2 to induce STAT5 phosphorylation provides direct evidence that a key functional output of IL-6 signaling has been conserved across approximately 500 million years of vertebrate evolution. Together with the presence of IL-1 (Fontenla-Iglesias et al., 2025) and IL-17 (Han et al., 2015) in lampreys, these findings point to a primordial inflammatory cytokine network that the emergence of somatically diversifying antigen receptors as a hallmark of vertebrate adaptive immunity.

Among the eight *SOCS* genes present in humans, *SOCS1*, *SOCS2*, *SOCS3*, and *CISH* are key regulators of cytokine signaling. In lamprey, we identified three *SOCS1/3*-like genes. In humans, SOCS1 inhibits JAK signaling downstream of IFN-γ, IFN-α/β, IL-12, IL-2, IL-7, IL-15, IL-21, and IL-4, whereas SOCS3 primarily restrains JAK signaling in response to IL-6, IL-23, and CSF3 (Sobah et al. 2021). In sea lamprey, two of the *SOCS1/3*-like genes were upregulated in response to IL-6.1 and IL-6.2 stimulation, revealing an ancient IL-6-SOCS regulatory axis that couples cytokine-induced STAT signaling to negative-feedback control.

Several aspects of IL-6 signaling in cyclostomes remain unresolved. Because we relied on cross-reactive anti-human STAT antibodies, we were able to monitor STAT5 but not STAT1 phosphorylation in sea lamprey. Moreover, our current experiments did not detect IL-6.3-induced STAT5 phosphorylation or SOCS induction, a result that may reflect engagement of a distinct downstream signaling axis and/or limitations of the recombinant IL-6.3 construct and expression system rather than a lack of intrinsic activity of the IL-6.3 paralog. Species-specific reagents, together with alternative IL-6.3 recombinant constructs and stimulation of additional cell and tissue targets, will be required to define the biological properties of IL-6.3 and to determine the conditions under which it may activate STAT signaling and induce *SOCS* genes. It will also be important to define the role of *IL-6* genes in the adaptive immune system, for example by testing their effects on antibody secretion by VLRB-expressing lamprey cells.

Collectively, our data demonstrate that jawless vertebrates possess a functional IL-6-STAT5 signaling axis that is inducible by a pathogen-associated stimulus and tissue injury and is coupled to SOCS-mediated negative feedback. By integrating structure-guided identification of *IL-6* orthologs with conserved genomic context, exon-intron organization, and paralog-specific patterns of leukocyte and tissue expression, we provide evidence that IL-6 forms part of an evolutionarily ancient inflammatory cytokine repertoire shared by jawed and jawless vertebrates. The ability of lamprey IL-6 paralogs to activate *STAT5* and *SOCS1/3*-like genes in myeloid cells, together with their distinct transcriptional responses to flagellin and tissue damage, offers a framework for dissecting how IL-6 cytokines were incorporated into vertebrate immune signaling networks. Broadly, this work illustrates how structure-guided comparative genomics and functional assays can resolve deeply divergent cytokine lineages and offers a framework for dissecting the emergence and diversification of vertebrate immune signaling pathways.

## Materials and Methods

### Animal husbandry

Non-feeding adult sea lampreys (45-50 cm) were obtained from the US Geological Survey Hammond Bay Biological Station and maintained in a 400-L tank at 16°C. Larval lampreys (8-15 cm) were acquired from local suppliers and housed in aquaria with sand substrate at 18°C. Prior to all experimental procedures, animals were anesthetized or euthanized with buffered MS-222 (Syndel) at final concentrations of 0.1 g/L or 1 g/L, respectively. All procedures were conducted in accordance with institutional and federal guidelines and were approved by the Emory University Institutional Animal Care and Use Committee (IACUC).

### Bioinformatics

Three-dimensional structural models of the proteins were generated using ESMfold (Lin et al. 2023) or AlphaFold (Jumper et al. 2021). Protein domains were annotated using the SMART web server (Letunic and Bork 2025) for domain prediction and structural similarity searches were conducted via the Foldseek Search Server (van Kempen et al. 2024). Predicted structures were visualized, aligned, and analyzed using PyMOL (Schrödinger, LLC).

Multiple sequence alignments were generated with T-Coffee using the PSI-Coffee mode (Notredame et al. 2000) and subsequently inspected by eye to verify alignment quality. Phylogenetic analyses were performed in MEGA (Saitou and Nei 1987; Tamura et al. 2021) version 12 using the maximum-likelihood method (Felsenstein 1981) with the JTT substitution model (Jones et al. 1992) and the pairwise deletion option. The reliability of the inferred topologies was evaluated by bootstrap resampling with 1,000 replicates, providing statistical support for the branching relationships. Accession numbers for all sequences included in the analyses are listed in Table S1.

Microsynteny was assessed by inspecting the 50 genes upstream and 50 genes downstream of the *IL-6* locus in selected vertebrate genomes using the NCBI Genome Data Viewer (https://www.ncbi.nlm.nih.gov/gdv). Analyses were performed on the following reference assemblies: *Homo sapiens* (GRCh38.p14), *Gallus gallus* (bGalGal1.mat.broiler.GRCg7b), *Lepisosteus oculatus* (fLepOcu1.hap2), *Amblyraja radiata* (sAmbRad1.1.pri), *Petromyzon marinus* (UKy_Petmar_22M1.pri1.0), and *Myxine glutinosa* (UKY_Mglu_1.0). Gene annotations were obtained from NCBI, and candidate orthologs were identified by reciprocal BLAST searche (Altschul et al. 1990). Conserved gene order was manually assessed to evaluate syntenic relationships and to infer evolutionary conservation of the *IL-6* genomic context.

Macrosynteny analysis was performed as previously described (Fontenla-Iglesias et al. 2025).Briefly, *Homo sapiens* (reference genome GRCh38.p14) and *Petromyzon marinus* (reference genome UKy_Petmar_22M1.pri1.0) unique orthologs were identified using OrthoFinder (Emms and Kelly 2019) and the resulting data were used as input to construct ribbon diagrams for selected species using the MacrosyntR package in R (El Hilali and Copley 2023). Significant associations were computed using Fisher’s exact test.

Prediction of phosphorylation sites in lamprey STAT proteins was performed using NetPhos 3.1.

### Flow cytometry and cell sorting

White blood cells were isolated from sea lamprey blood for analysis by immunofluorescence flow cytometry following the protocol described previously (Guo et al. 2009; Hirano et al. 2013). Briefly, Percoll-separated leukocytes from lamprey blood were stained with specific antibodies, including mouse anti-VLRB (4C4), anti-VLRC (3A5), anti-VLRA (9A3), anti-monocyte (8A1), and anti-granulocyte (2D4) monoclonal antibodies. Cells were gated based on forward scatter-A (FSC-A) vs. side scatter-A (SSC-A) for leukocyte populations, FSC-A vs. FSC-H for singlets, and exclusion of dead cells using negative LIVE/DEAD Aqua (Invitrogen) staining. Flow cytometric analyses were performed using a MACSQuant Analyzer (Miltenyi Biotec). VLRA+, VLRB+, VLRC+, and VLR triple-negative (TN) cells, as well as monocytes and granulocytes, were sorted using a BD FACS Aria II (BD Biosciences) for subsequent qPCR analysis. The purity of sorted populations exceeded 90%.

### Experimental immune stimulation protocols

Unstimulated lamprey ammocoetes (n=3 per group) were injected intraperitoneally with 20 µL of *Salmonella typhimurium* flagellin (50 ng/g body weight; Millipore Sigma) diluted in 0.66× PBS, or with 20 µL of 0.66× PBS alone as a control. Typhlosole/ intestine tissues were collected at 2, 8, and 24 hours post-injection for reverse transcription quantitative PCR (RT-qPCR) analysis.

Alternatively, naïve lamprey ammocoetes (n=6 per group) were maintained overnight in autoclaved water supplemented with penicillin/streptomycin. Animals were then mechanically wounded by puncturing the skin with a sterile needle. Negative controls were handled identically but not wounded. Two hours post-injury, animals were euthanized and bled, and ∼0.2 cm² of skin surrounding the wound site was dissected and preserved for RT-qPCR analysis.

To isolate peritoneal exudate cells, we adapted a murine model previously described (Stewart et al. 1975). Briefly, non-feeding adult sea lampreys were injected intraperitoneally with 2 mL of 4% thioglycolate solution prepared in water, autoclaved, and aged for 3 months in the dark at room temperature. After 3 days, animals were euthanized and bled, and the peritoneal cavity was washed with sterile 0.66× PBS to collect exudate cells. The recovered cells were washed three times in 0.66× PBS, then kept on ice for immediate downstream *in vitro* applications or snap-frozen in dry ice for storage.

### RNA-seq data processing and quantification

Raw sequencing reads were evaluated for quality using FastQC (v0.12.1). Adapter sequences and low-quality bases were removed with Trimmomatic (v0.40) (Bolger et al. 2014) using default Illumina parameters. The filtered reads were quantified with Salmon (v1.10) (Patro et al. 2017), using the current genome reference annotation (UKy_Petmar_22M1.pri1.0). Transcript-level quantifications were imported into R and summarized to the gene level to obtain transcripts-per-million (TPM) values, which were used for downstream expression analyses.

### *In vitro* Stimulation of Peritoneal Exudate Cells

Peritoneal exudate cells (5 × 10⁵ viable cells per well) were resuspended in 0.66× PBS supplemented with 0.5% fetal bovine serum (FBS) and plated in 24-well plates. Cells were stimulated at 16 °C for 0.5, 1, or 3 hours with 200 ng/mL of recombinant IL-6.1, IL-6.2, or IL-6.3 proteins, or with 20 ng/mL of *Salmonella typhimurium* flagellin. Following stimulation, cells were centrifuged at 500 × g for 5 minutes at 4 °C and lysed directly in the wells by adding RLT buffer (Qiagen) for downstream RNA extraction.

### Quantitative Real-Time PCR

Total RNA from lamprey tissues or cell populations was extracted using the RNeasy Mini Kit (Qiagen), including on-column DNase digestion to remove genomic DNA. First-strand cDNA was synthesized with the LunaScript RT SuperMix Kit (New England Biolabs). Quantitative real-time PCR was carried out using SYBR Green dye on a 7900HT ABI Prism system (Applied Biosystems). Relative gene expression was normalized to β-actin. All primers used in this study are listed in Supplementary Table 2.

### Immunoblot

Peritoneal exudate cells were plated at a density of 2 × 10⁶ cells per well in 12-well plates and stimulated as described above. At 15, 30, and 60 minutes post-stimulation, cells were lysed directly in NP-40 lysis buffer (Thermo Fisher) supplemented with protease inhibitor cocktail (Roche) and activated sodium orthovanadate (Na₃VO₄; 1 mM) (Sigma-Aldrich). Protein concentration was quantified by a NanoDrop spectrophotometer, and 30 µg of total reduced protein per sample was run on a 4-20% precast SDS-PAGE gel (Bio-Rad) and transferred to a PVDF membrane (Millipore) using a semi-dry transfer system. Membranes were blocked in 5% skim milk in TBS-T (TBS with 0.1% Tween-20) for 1 hour at room temperature and incubated overnight at 4 °C with primary antibodies anti-STAT1 Phospho (Tyr701) (Biolegend), anti-STAT5 Phospho (Tyr694) (Biolegend) or anti-β-ACTIN (Biolegend). After washing, membranes were incubated with HRP-conjugated secondary anti-mouse antibodies (BioLegend) for 1 hour at room temperature. Detection was performed using enhanced chemiluminescence (ECL) reagents (Thermo Fisher) and imaged with a ChemiDoc Imaging System (Bio-Rad). To validate the phospho-specific antibodies, peritoneal exudate cells were treated with pervanadate to induce global tyrosine phosphorylation, and an anti-phosphotyrosine antibody was included as a positive control.

## Supporting information

IL-6-manuscript-04-02-2026.pdf

## Acknowledgements

This study was supported by National Institutes of Health grants R01AI072435, R35GM122591, and GM108838; the Georgia Research Alliance; the Max Planck Society; the European Research Council (ERC) under the European Union’s Seventh Framework Program (FP7/2007-2013), grant agreement 323126; PID2020-113087RB-I00 from the Ministry of Science and Innovation and the European Regional Development Fund (FEDER, European Union); and grant ED431C2017/31 from the Xunta de Galicia. We thank Mr. R.E. Karaffa II and K. Fife from the Emory University School of Medicine Flow Cytometry Core for assistance with cell sorting. Bulk RNA-sequencing services were provided by the Emory NPRC Genomics Core, which is supported in part by NIH P51 OD011132. Several figures were created with BioRender.com.

## Author Contributions

F.F-I and S.D. contributed to the study design. F.F-I., S.D., J.B., J.P.R., M.H., B.B.A., J.L., W.L., T.B., and M.D.C. performed the experiments, and/or analyzed the data, and supervised the study. S.D., F.F-I., T.B., J.P.R., and M.D.C. wrote the manuscript. All authors reviewed the manuscript and provided feedback.

