## Supplementary material for "Ancient IL-6-STAT5 signaling orchestrates inflammation in jawless vertebrates": IL-6-manuscript-04-02-2026.pdf

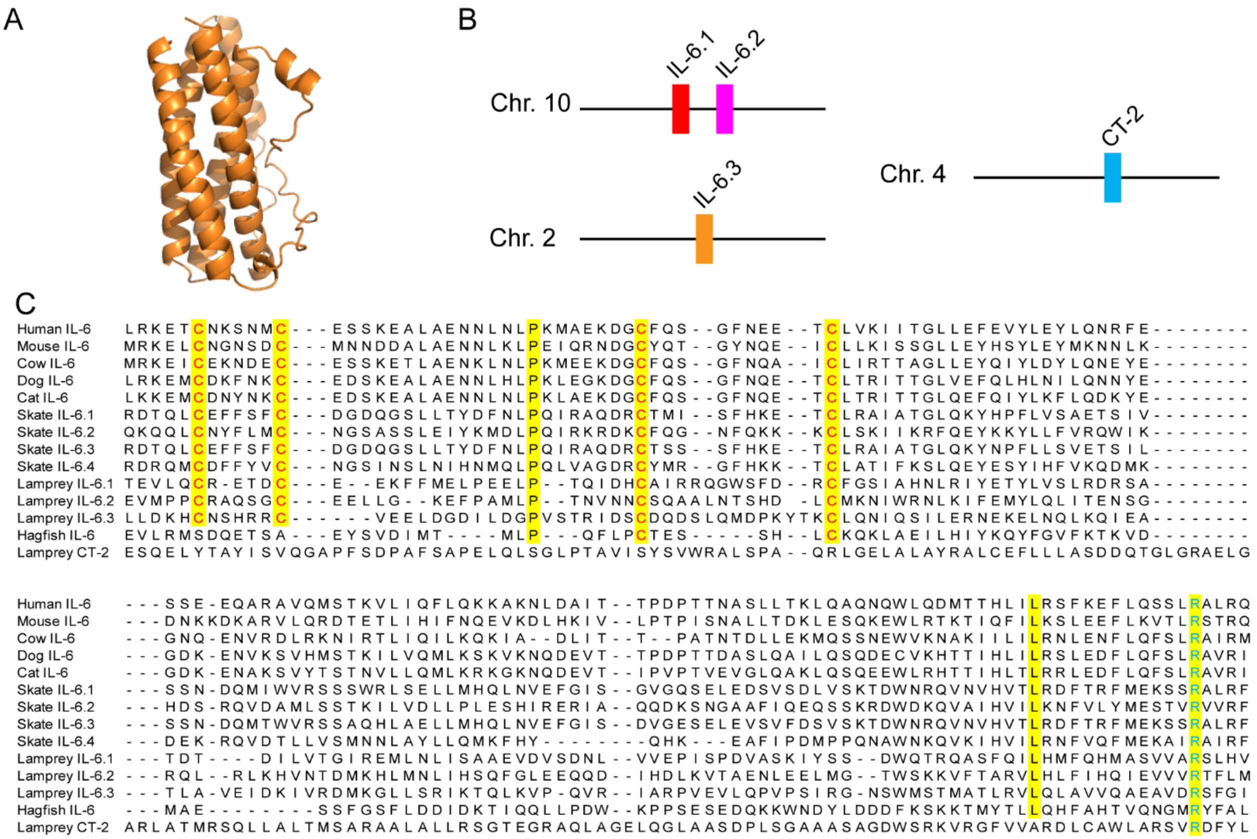


**Supplementary Figure 1.** Structural, genomic, and sequence relationship between lamprey CT-2-like and IL-6-related cytokines. (A) Predicted three-dimensional structure of the sea lamprey CT-2-like protein, showing the characteristic four-helix bundle also observed in other IL-6 family cytokines (see Fig. 1). (B) Genomic distribution of the three lamprey IL-6-like genes and the CT-2-like gene. IL-6.1 and IL-6.2 map to chromosome 10, IL-6.3 to chromosome 2, and CT-2-like to chromosome 4. (C) Multiple sequence alignment of IL-6 proteins from representative jawed vertebrates with candidate IL-6-like sequences identified in jawless vertebrates. The four canonical cysteine residues of vertebrate IL-6, together with additional conserved positions, are highlighted. The hagfish IL-6 candidate retains only two of these four cysteines, whereas the lamprey CT-2-like sequence lacks conservation at the corresponding sites.

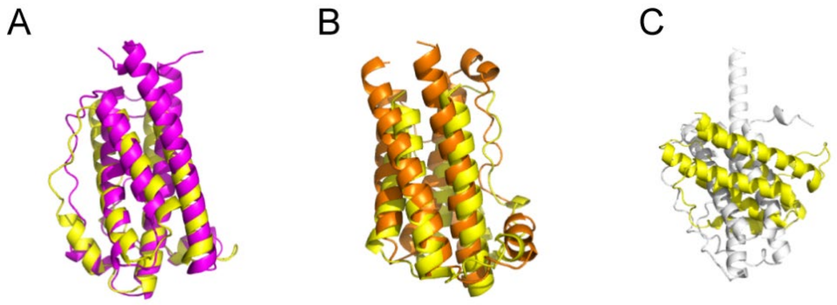


**Supplementary Figure 2.** Structural alignments of the three sea lamprey IL-6 sequences (IL-6.1, IL-6.2, and IL-6.3) with hagfish IL-6 (yellow). IL-6.1 and IL-6.2 display strong topological conservation relative to hagfish IL-6 (A, B), whereas IL-6.3 exhibits weaker structural conservation (C).

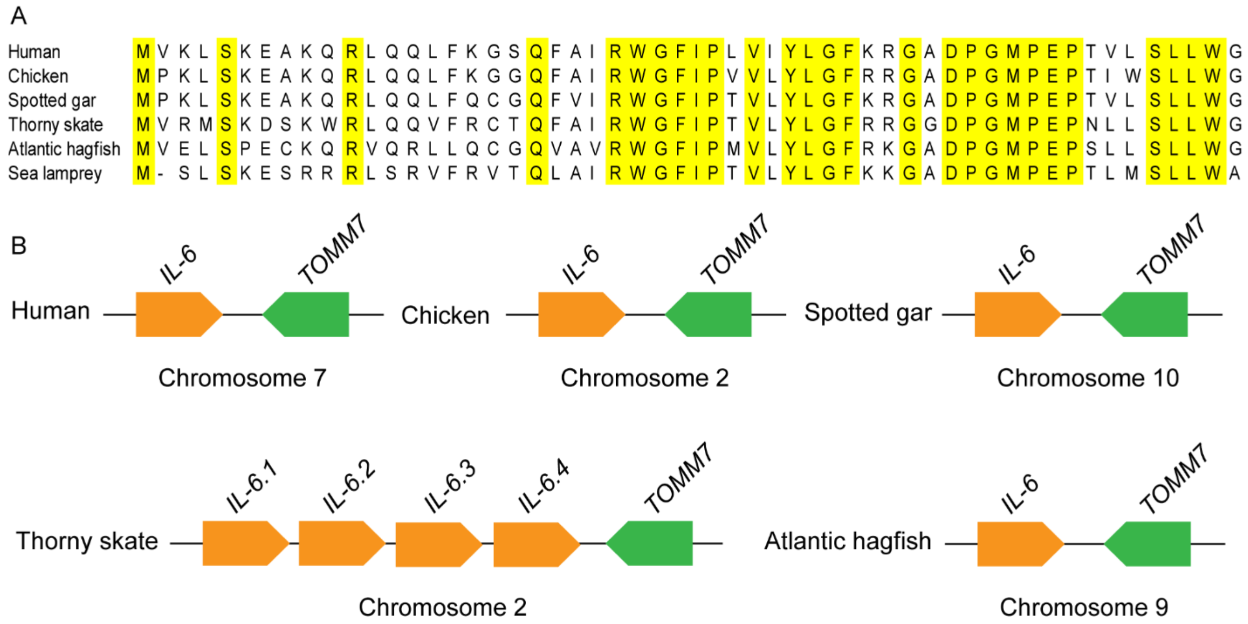


**Supplementary Figure 3.** *TOMM7* (translocase of the outer mitochondrial membrane 7) and its genomic relationship with *IL-6*. (A) Multiple sequence alignment of translated TOMM7 proteins. Conserved amino acid residues are highlighted in yellow. (B) Genomic organization of *IL-6* and *TOMM7*, showing that the two genes are positioned adjacent to one another and transcribed in opposite orientations. In lamprey, however, *TOMM7* is located at a different genomic location (see Fig. 2). The schematic is not drawn to scale.

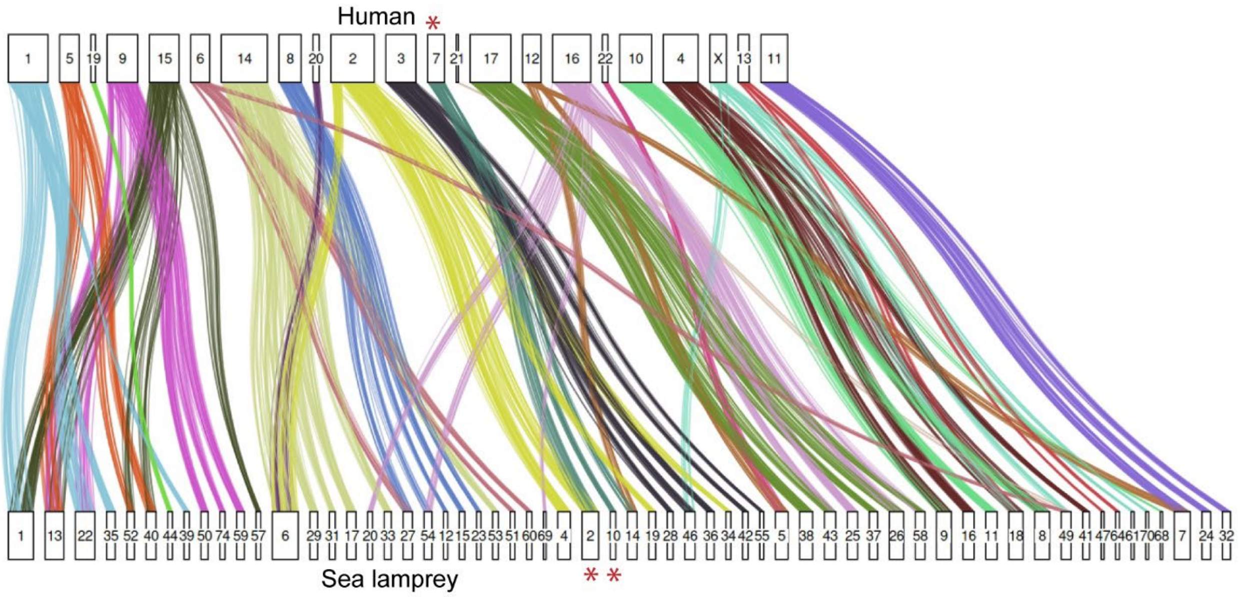


**Supplementary Figure 4.** Chromosomal macrosynteny between human and sea lamprey. Asterisks indicate chromosomes harboring IL-6 loci. Enrichment of shared orthologs between these chromosomes ($p \leq 0.05$, Bonferroni-corrected one-sided Fisher's exact test) supports an ancestral relationship between the *IL-6*-containing chromosomal segments. Each box denotes a chromosome, and vertical connectors represent orthologous gene pairs.

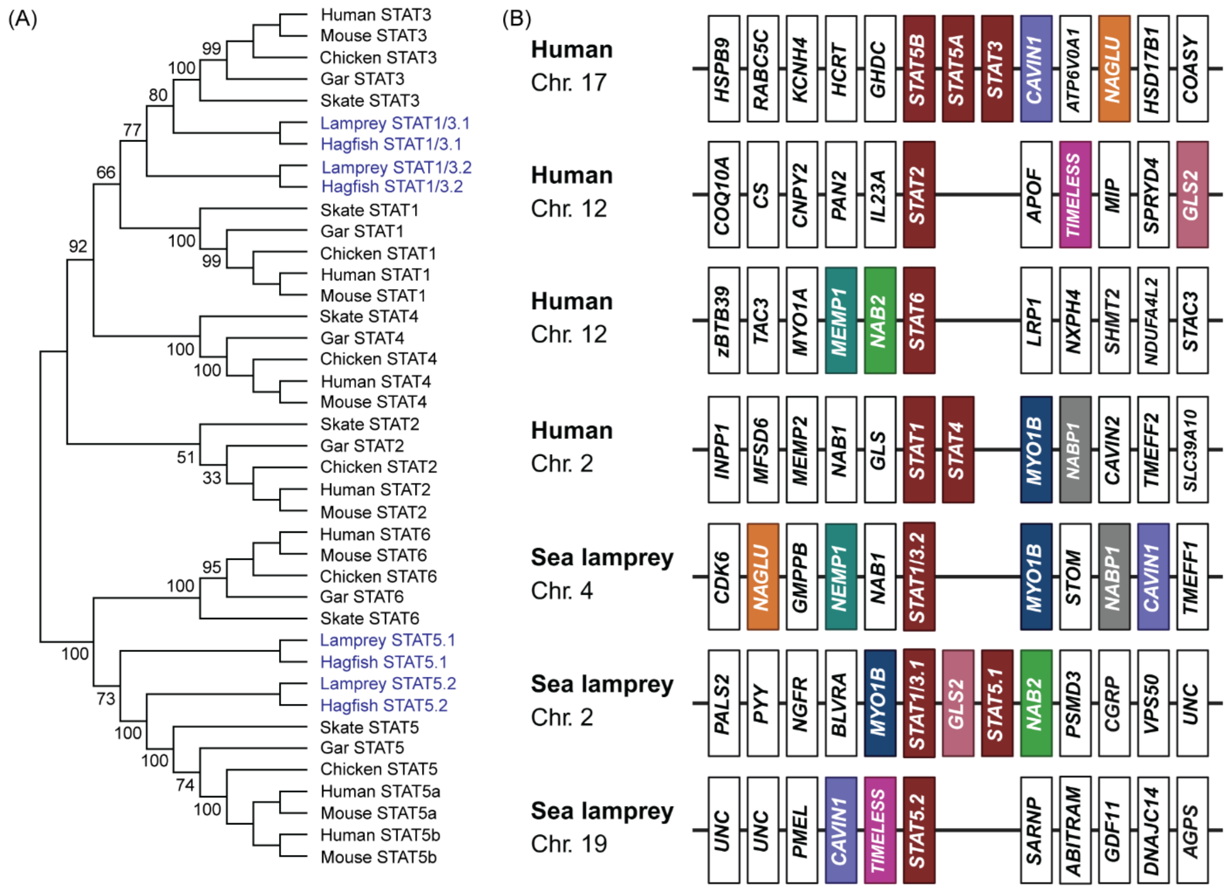


**Supplementary Figure 5.** Phylogenetic and syntenic analysis of STAT family genes in jawless vertebrates. (A) Maximum-likelihood phylogenetic analysis of STAT family proteins from representative jawed vertebrates and homologs identified in lamprey and hagfish. Two agnathan homolog groups, STAT1/3.1 and STAT1/3.2, cluster near gnathostome STAT1 and STAT3, whereas STAT5.1 and STAT5.2 group with gnathostome STAT5. Agnathan sequences are shown in blue, and numbers at nodes indicate bootstrap support values. (B) Conserved synteny analysis of genomic regions spanning the five upstream and five downstream flanking genes surrounding human and sea lamprey STAT loci. Human STAT loci on chromosomes 17, 12, and 2 exhibit a mosaic pattern of syntenic conservation with three sea lamprey STAT-related regions. The lamprey chromosome 4 locus, containing STAT1/3.2, retains NAGLU and CAVIN1, consistent with conservation relative to the human chromosome 17 locus, and also shares MYO1B, NABP1, and NEMP1 with human loci on chromosomes 2 and 12. The lamprey chromosome 2 region, containing STAT1/3.1 and STAT5.1, shares MYO1B with human chromosome 2 and GLS2 and NAB2 with human chromosome 12. The lamprey chromosome 19 locus, containing STAT5.2, retains CAVIN1 and TIMELESS, further linking it to the human chromosome 17 and 12 STAT regions. Colored boxes indicate conserved syntenic genes; white boxes denote neighboring genes not identified as conserved. Genomic regions are shown schematically and are not to scale.

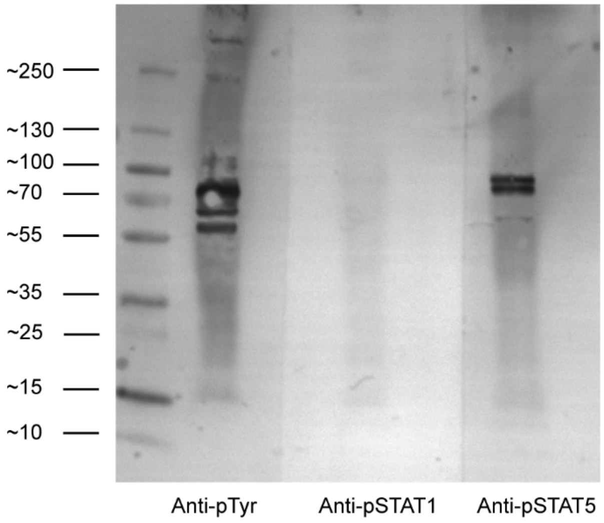


**Supplementary Figure 6.** Validation of phospho-STAT antibodies in lamprey peritoneal exudate cells. Immunoblot analysis of reduced protein lysates from pervanadate-treated lamprey peritoneal exudate cells. Lane 1, molecular weight marker. Membranes were probed with anti-phosphotyrosine, anti-phospho-STAT1, and anti-phospho-STAT5 antibodies. The anti-phosphotyrosine antibody detected strong phosphorylation signals, confirming effective induction of tyrosine phosphorylation by pervanadate. Under these conditions, the anti-phospho-STAT5 antibody showed detectable reactivity, whereas the anti-phospho-STAT1 antibody did not yield a detectable signal. These results support the use of the phospho-STAT5 antibody, but not the phospho-STAT1 antibody, for analyses in lamprey peritoneal exudate cells.

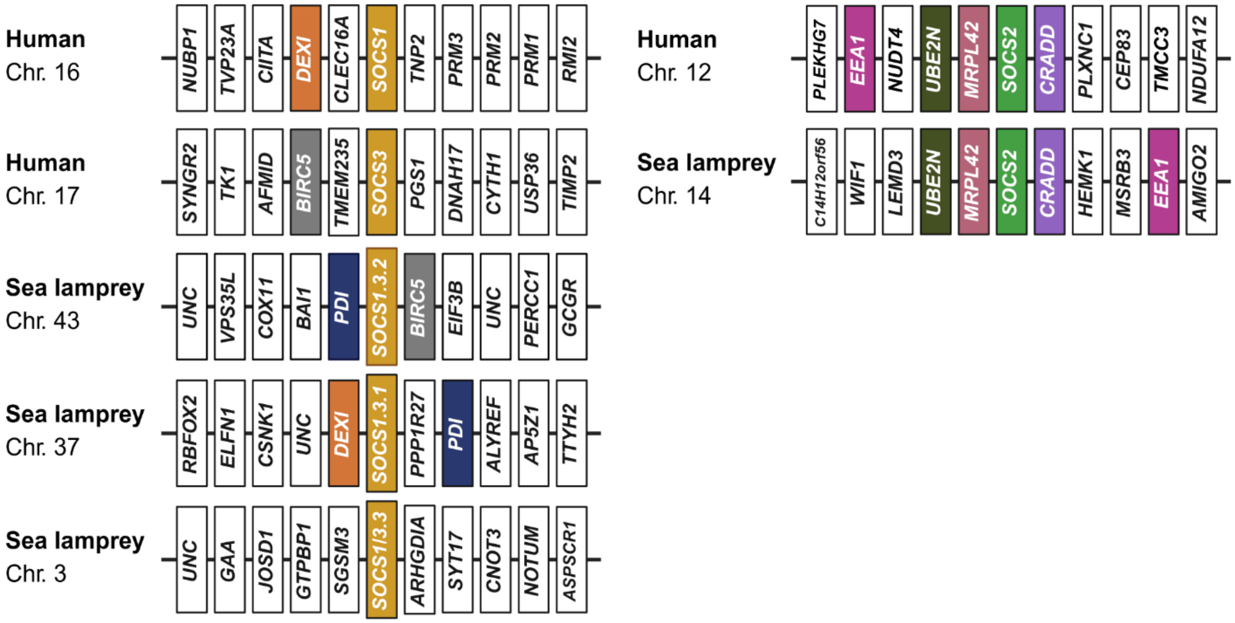


**Supplementary Figure 7.** Conserved synteny analysis of SOCS family loci in human and sea lamprey. Genomic regions encompassing the five upstream and five downstream flanking genes surrounding members of the SOCS1–3 subfamilies were compared between human and sea lamprey. Limited local syntenic conservation was detected between the human SOCS1 and SOCS3 loci and two sea lamprey SOCS1/3-related loci. Specifically, the SOCS1/3.1 locus on chromosome 37 retains DEXI, whereas the SOCS1/3.2 locus on chromosome 43 retains BIRC5, indicating residual conservation of the genomic neighborhoods associated with the human SOCS1 and SOCS3 loci, respectively. Both lamprey loci also contain PDI, further supporting a relationship between these two chromosomal segments. By contrast, the lamprey SOCS1/3.3 locus on chromosome 3 does not exhibit clear local synteny with either human SOCS1 or SOCS3. In comparison, the SOCS2 locus displays stronger local syntenic conservation between human chromosome 12 and sea lamprey chromosome 14, including shared flanking genes such as UBE2N, MRPL42, and CRADD. Colored boxes indicate conserved syntenic genes, whereas white boxes denote neighboring genes not identified as conserved. Genomic regions are shown schematically and are not drawn to scale.

**Supplementary Table 1.** Sequences used for phylogenetic and comparative analyses.

| Organism | Sequence | Genomic location and/or accession number |
|---|---|---|
| Sea lamprey (kPetMar1.pri) | IL-6.1 | Ch.10:11054250-11060242:F, XP_032807189, XM_032951298 |
| | IL-6.2 | Ch.10:11040338-11046597:F, XP_032807190, XM_032951299 |
| | IL-6.3 | Ch.2:10651754-10656909:R, XP_032819639, XM_032963748 |
| | TOMM7 | Sc.443:39653-50055:F, XP_032836492, XM_032980601 |
| | CT2 | Ch.4:1057793-1064544:R, XP_032802522, XM_032946631 |
| | STAT1/3.1 | XP_032814978 |
| | STAT1/3.2 | XP_032802147 |
| | STAT5.1 | XP_075927610 |
| | STAT5.2 | XP_032813201 |
| | SOCS1/3.1 | XP_032823113, XM_032967222 |
| | SOCS1/3.2 | XP_032826112, XM_032970221 |
| | SOCS1/3.3 | XP_032834399, XM_032978508 |
| | SOCS2 | XP_032810244, XM_032954353 |
| Pouched lamprey (kcGeoAust1.1) | IL-6.1 | Sc.hic_geoau00000006:4225354-4229819:F |
| | IL-6.2 | Sc.hic_geoau00000006:4210276-4215810:F |
| | IL-6.3 | Sc.hic_geoau00000005:14009500-14013636 |
| | TOMM7 | Sc.hic_geoau00000003:6873391-6874263:R |
| Atlantic hagfish (UKY_Mglu_1.0) | IL-6 | Ch.9:93258752-93262676:R, XP_067981902, XM_068125801 |
| | TOMM7 | Ch.9:93204505-93257059:F, XP_067983059, XM_068126958 |
| | STAT1/3.1 | XP_067975739 |
| | STAT1/3.2 | XP_067979438 |
| | STAT5.1 | XP_067954492 |
| | STAT5.2 | XP_067984644 |
| | SOCS1/3.1 | XP_067972766 |
| | SOCS1/3.2 | XP_067973305 |
| | SOCS1/3.3 | XP_067966582 |
| | SOCS2 | XP_067968758 |
| Thorny skate (sAmbRad1.1.pri) | IL-6.1 | Ch.2:51668239-51669628:F, XP_032894266, XM_033038375 |
| | IL-6.2 | Ch.2:51707818-51712158:F, XP_032902107, XM_033046216 |
| | IL-6.3 | Ch.2:51726132-51730972:F, XP_032902121, XM_033046230 |
| | IL-6.4 | Ch.2:51784335-51800348:F, XP_032895270, XM_033039379 |
| | TOMM7 | Ch.2:51811311-51838923:R, XP_032902132, XM_033046241 |
| | STAT1 | XP_032879642 |
| | STAT2 | XP_032871486 |

| | | |
|---|---|---|
| | STAT3 | XP_032891109 |
| | STAT4 | XP_032879641 |
| | STAT5 | XP_032891107 |
| | STAT6 | XP_032871482 |
| | SOCS1 | XP_032896294, XP_032869201 |
| | SOCS2 | XP_032895307, XP_032898399 |
| | SOCS3 | XP_032900159 |
| | CISH | XP_032892912 |
| Spotted gar (LepOcu1) | IL-6 | Ch.10:1358682-1361152:R, XP_015213289, XM_015357803 |
| | TOMM7 | Ch.10:1343960-1355035:F, XP_006636264, XM_006636201 |
| | STAT1 | XP_015214635 |
| | STAT2 | XP_006629507 |
| | STAT3 | XP_015217376 |
| | STAT4 | XP_015214803 |
| | STAT5 | XP_015217370 |
| | STAT6 | XP_069043157 |
| | SOCS1 | XP_006637341 |
| | SOCS2 | XP_015207859, XP_006628404 |
| | SOCS3 | XP_015212237 |
| | CISH | XP_069045716 |
| Chicken (GRCg6a) | IL-6 | Ch.2:30863330-30865609:F, NP_989959, NM_204628 |
| | TOMM7 | Ch.2:30874543-30877963:R, NP_001186546, NM_001199617 |
| | STAT1 | NP_001012932 |
| | STAT2 | XP_040510813 |
| | STAT3 | NP_001385253 |
| | STAT4 | XP_015144934 |
| | STAT5 | XP_046760288 |
| | STAT6 | XP_046760984 |
| | SOCS1 | NP_001131120 |
| | SOCS2 | NP_989871 |
| | SOCS3 | NP_001131120 |
| | CISH | NP_989957 |
| Human (GRCh38.p14) | IL-6 | Ch.7:22727521-22731567:F, NP_000591, NM_000600 |
| | TOMM7 | Ch.7:22822779-22818002:R, NP_061932, NM_019059 |
| | IL-11 | NP_000632 |
| | IL-31 | NP_001014358 |
| | OSM | NP_065391 |
| | LIF | AAA51699 |
| | CLCF1 | NP_037378 |
| | CNTF | NP_000605 |
| | CT1 | NP_001321 |
| | STAT1 | NP_001371819 |
| | STAT2 | NP_001372039 |

| | | |
|---|---|---|
| | STAT3 | NP_001371918 |
| | STAT4 | NP_001230764 |
| | STAT5 | NP_001275647, NP_036580 |
| | STAT6 | NP_001171549 |
| | SOCS1 | NP_003736 |
| | SOCS2 | NP_003868 |
| | SOCS3 | NP_003946 |
| | CISH | NP_037456 |
| Mouse (GRCm39) | IL-6 | Ch.5:30218190-30224546:F, NP_112445, NM_031168 |
| | IL-11 | NP_032376 |
| | IL-31 | NP_083870 |
| | OSM | NP_065391 |
| | LIF | NP_032527 |
| | CLCF1 | NP_064336 |
| | CNTF | NP_000605 |
| | CT1 | NP_031821 |
| | CT2 | NP_942155 |
| | STAT1 | NP_033309 |
| | STAT2 | NP_064347 |
| | STAT3 | NP_998825 |
| | STAT4 | NP_001295195 |
| | STAT5 | NP_035618, NP_001107035 |
| | STAT6 | NP_033310 |
| | SOCS1 | NP_034026 |
| | SOCS2 | NP_031732 |
| | SOCS3 | NP_031733 |
| | CISH | NP_034025 |
| Cow | IL-6 | NP_776348 |
| | IL-11 | XP_024835139 |
| | IL-31 | ELR49822 |
| | OSM | NP_783644 |
| | LIF | AAC27535 |
| | CLCF1 | XP_024843329 |
| | CNTF | XP_003587079 |
| | CT1 | NP_001179313 |
| | CT2 | XP_070218673 |
| Dog | IL-6 | NP_001003301 |
| | IL-11 | XP_022283255 |
| | IL-31 | NP_001159386 |
| | OSM | XP_005636497 |
| | LIF | NP_001184002 |
| | CLCF1 | XP_025302004 |
| | CNTF | XP_005631242 |
| | CT1 | NP_001274070 |
| | CT2 | XP_022276378 |
| Cat | IL-6 | NP_001009211 |
| | IL-11 | XP_023101181 |

| | | |
|---|---|---|
| | IL-31 | XP_044897990 |
| | OSM | XP_044897937 |
| | LIF | XP_003994833 |
| | CLCF1 | XP_023095876 |
| | CNTF | XP_003993451 |
| | CT1 | XP_011288837 |
| | CT2 | XP_044904219 |

Note: Genome assembly versions are indicated in parentheses for the organisms included in the genomic analysis. In humans, the CT-2 locus is annotated as a pseudogene and was therefore excluded from this analysis.

**Table S2.** Primers used for gene expression analysis by RT-PCR in this study.

| Primer name | Sequence (5'-3') | Purpose |
|---|---|---|
| qIL6.1-F | ACATAACGGAGGTGCTGCAA | qPCR |
| qIL6.1-R | AAACTCCACCCCTGTCTCCT | qPCR |
| qIL6.2-F | AACACCGACATGAAGCACCT | qPCR |
| qIL6.2-R | CCACGTTCCCATCAGCTCTT | qPCR |
| qIL6.3-F | TGAGCCCTCTATTCCGGTGT | qPCR |
| qIL6.3-R | CATCCAGCTCCTCCACACAG | qPCR |
| qSOCS1/3.1-F | CGTATCGGGATTAGAGCGGAC | qPCR |
| qSOCS1/3.1-R | GACCCGGGCAGATGAACC | qPCR |
| qSOCS1/3.2-F | TCGAGTGCATCATCAGCCTG | qPCR |
| qSOCS1/3.2-R | CTTCCTCCTCCTCCTCCTCC | qPCR |
| qIL8-F | CCATCCCAAGCATTTCCAGACA | qPCR |
| qIL8-R | GGTGTCTGAGCCCCGTCCAAAA | qPCR |
| qbActin-F1 | GCCAACCGTGAAAAGATGACA | qPCR |
| qbActin-R1 | GGATGGCGACGTACATTGC | qPCR |
